## Supplementary figures for "Versatile live-cell activity analysis platform for characterization of neuronal dynamics at single-cell and network level"

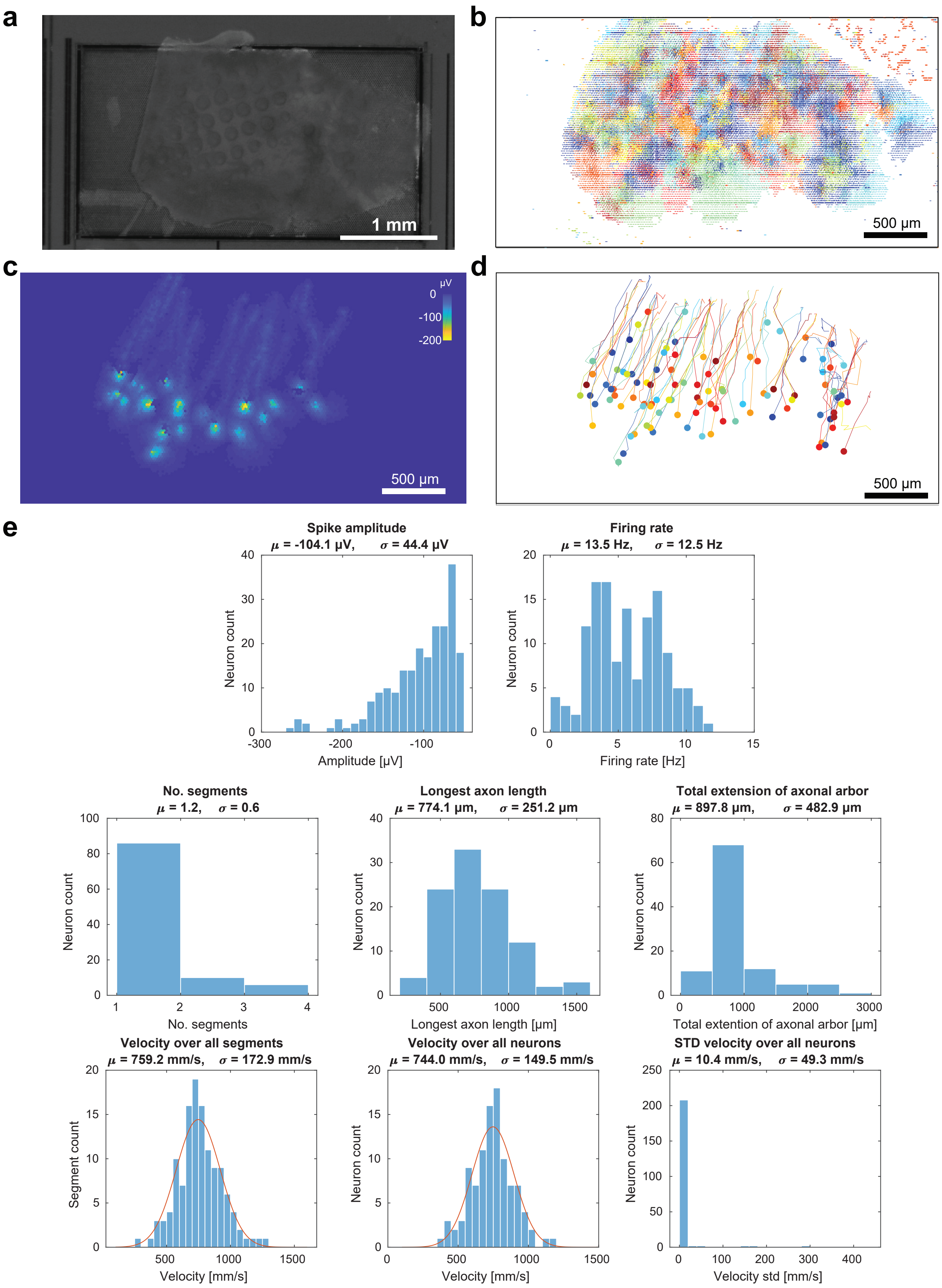


Suppl. Figure. 1: Extraction of axonal signal propagation characteristics from the retina recordings. (a) Microscopy image of a retina placed on the DM-MEA. (b) Extracted electrical footprints for n=202 spike-sorted neurons, showing a complete coverage of the retina. (c) Example footprints of 20 neurons. (d) Extracted axonal segments for all detected neurons, showing one axon originating from each neuron. All axons are oriented in the same direction. (e) Statistics extracted for the retina (n = 202 neurons) with the same methods that have been used to obtain the panels in Fig 3, including spike amplitude and firing rate for each neuron, axon lengths and propagation velocities. Mean values (μ) are given as well as standard deviations (σ).

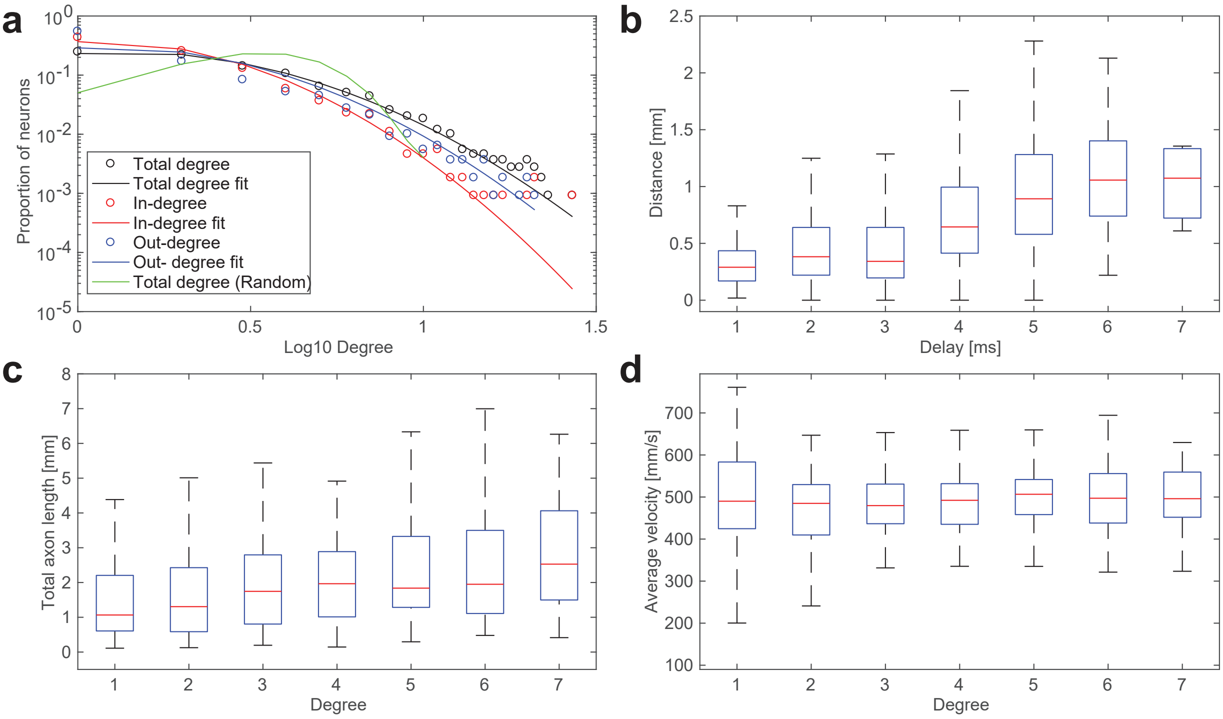


Suppl. Figure. 2: Analysis of network connectivity, and correlation between single-cell properties and network connectivity. (a) Degree distribution (in-degree, out-degree and total degree) with log-normal fitting and comparison to random directional network. (b) Correlation between connection length and time delay. (c) Correlation between total degree and total extension of axonal arbor. (d) Correlation between total degree and average velocity.
